# Brain-wide transcriptome-based metabolic alterations in Parkinson’s disease: human inter-region and human-experimental model correlations

**DOI:** 10.1101/2022.08.31.505965

**Authors:** Regan Odongo, Orhan Bellur, Ecehan Abdik, Tunahan Çakır

## Abstract

Alterations in brain metabolism are closely associated with the molecular hallmarks of Parkinson’s disease (PD). A clear understanding of the main metabolic perturbations in PD is therefore important. Here, we retrospectively analysed the expression of metabolic genes from 34 PD-control post-mortem human brain transcriptome data from literature, spanning multiple brain regions, and found significant metabolic correlations between the Substantia nigra (SN) and cerebral cortical tissues with high perturbations in protein modification, transport, nucleotide and inositol phosphate metabolic pathways. Moreover, three main metabolic clusters of SN tissues were identified from patient cohort studies, each characterised by perturbations in (a) pyruvate, amino acid, neurotransmitter, and complex lipid metabolisms (b) inflammation-related metabolism, and (c) lipid breakdown for energy metabolism. Finally, we analysed 58 PD-control transcriptome data from in vivo/in vitro disease models and identified experimental PD models with significant correlations to matched human brain regions. Collectively, our findings are based on 47 PD transcriptome datasets covering 92 PD-control comparisons spanning more than 1000 samples in total, and they suggest metabolic alterations in several brain regions, heterogeneity in metabolic alterations between study cohorts for the SN tissues and suggest the need to optimize current experimental models to advance research on metabolic aspects of PD.

## Introduction

Altered metabolism is a major molecular event in most neurodegenerative diseases ^1,2^ including PD. Given that the main symptoms of this disease, motor and non-motor impairments, are associated with the functional coordination between multiple brain regions and that clinical symptoms vary among patients, metabolic alterations and their roles in PD might be the essential components of a new paradigm for identifying diagnostic metabolic biomarkers, paving the way for the discovery of novel drug targets.

The human brain is the second-most metabolically active organ in the body, and several studies using high throughput omics data, molecular biology and radiological techniques have provided lines of evidence for altered metabolite concentrations between healthy controls and PD ^3^. Like other neurodegenerative disorders, disturbances of the protein modifying and clearing systems, termed the ubiquitin-proteosome system, are strongly linked with PD pathology ^4–6^. Additionally, pathways related to energy generation through oxidative phosphorylation is frequently dysregulated in PD and are associated with hallmarks of the disease through mitochondrial dysfunction ^4,7–9^. Recent findings have suggested that lipid metabolism may play a major role in PD than previously thought as alterations in major brain lipids such as glycolipids, phosphatidylcholine, phosphatidylethanolamine, phosphatidylinositol and phosphatidylserine ^1,5,10,11^ are correlated with a decrease in brain mass, pathological inflammation and cell membrane integrity perturbations. Perturbations in metabolite transport reactions and neurotransmitter metabolism have also been reported ^4,7^. However, a comprehensive understanding of the metabolic changes including the main metabolic pathways in different brain regions affected during PD and their association with disease symptoms is yet to be fully explored.

Lewy body (LB) deposits are the main pathology associated with PD and its comorbidity with neuropsychological disorders are diagnostic molecular biomarkers routinely used to definitively distinguish PD from other neurodegenerative disorders ^4,12,13^. While LBs are almost frequently highly deposited in the substantia nigra (SN), Braak et al. ^12^ showed that pathologic aggregated alpha-synuclein (α-Syn), the main constituent of LB, follows a trajectory traceable from the gut in the affected individuals to the cortical regions of the brain following a prion-like propagation pattern ^12,14^. However, how the migrating α-Syn and resultant LB deposits alter metabolism in the affected brain regions and the resultant effects on the basal ganglia-cortex-cerebellum (BG-CX-CE) molecular circuitry implicated in PD ^13^ have been hardly investigated.

With accumulating high throughput omics data, systems biology methods offer holistic approaches for molecular pattern identification in disease biomarker discovery. To date, several methods have been applied on transcriptome, genome, proteome, and metabolome data to identify distinguishing molecular features underlying PD pathology ^3,15,16^. However, while these methods are easy to implement and interpret, they are less robust in applications involving high-dimensional datasets and, hence, hybrid approaches combining various techniques in statistics and machine learning are ideal for comprehensive elucidation of the underlying disease biology. Within this context, recently, Li and colleagues ^17^ combined metabolic gene information from Recon 3D ^18^ with transcriptome data from PD, Alzheimer’s and Huntington disease and identified metabolic differences and common disease biomarkers. However, this study was not PD-centred, and only one dataset with limited brain region coverage was analyzed, thereby potentially occluding our complete understanding of metabolism in PD. Thus, a systematic analysis leveraging molecular changes captured by different PD study cohorts is necessary to provide a comprehensive characterization of the metabolic alterations and evaluate potential mirroring experimental animal models.

In the current work, we performed a systematic analysis to unravel the metabolic landscape during PD from human brain post-mortem transcriptome data. We show the existence of (i) three main PD patient clusters with cluster-specific metabolic alterations in the SN, and (ii) a high correlation between gene expression levels in SN and cortical regions than other brain regions during PD. Finally, we analysed PD experimental models and prioritized those with high correlation with corresponding human brain region tissues as potentially useful for metabolic modelling of PD in future studies. Our study reaffirms the importance of brain metabolism during PD and demonstrates associated regional metabolic differences. We also conclude that not all available experimental models can be used to simulate altered metabolism in the human brain and thus PD pathology might be a function of systemic failure, which might be irreducible to single genetic/chemical perturbations as applied during the generation of most of the available experimental models in the literature.

## Results

### Gene expression data collection and pre-processing

Transcriptome datasets from post-mortem human brain tissues and from experimental models (cell line, mouse and rat) used in this study covered (i) the midbrain (basal ganglia including putamen (PUT), globus pallidus (GP), striatum (ST) and substantia nigra (SN)), amygdala (AM), hindbrain (CE), (ii) brainstem (medulla oblangata (ME), locus coeruleus (LC), dorsal motor nucleus of the vagus (DON) and inferior olivary nucleus (ION)) and (iii) the cerebral cortex (prefrontal cortex (PCX), whole cortex (CX), medial temporal (MTG) and superior frontal gyrus (SFG)) regions (detailed information in **Supplement 2 (Table S1**). These brain regions are amongst the most affected brain tissues in PD pathology with known functional and structural connections and have been implicated in PD-associated key phenotypes ^12,13,32– 34^. Additionally, sporadic PD cases were overrepresented in the collected datasets as compared to familial PD cases. Overall, our study represents one of the most comprehensive PD transcriptome data analysis based on metabolic genes available to date as compared to the most recent analysis ^17^, covering 92 comparisons from 47 post-mortem/in vivo model/in vitro model transcriptome datasets. Moreover, these datasets were composed of patient cohorts from different regions/countries allowing for cross-cohort comparison to identify those with similar metabolic profiles. The gene score ^30^ approach, which uses information regarding the specific gene fold change and corresponding p-value, was applied here. The unsupervised PCA-based filtering ^28^ based on gene scores was used to identify genes with highest variability across the comparisons, leading to a representative panel of 1525 genes from the imputation pre-processed 1833 genes when all the human brain tissues were used as input, and 1103 genes from 1521 genes when only data from SN region was used. Overall, in this gene panel, the most imputed gene had a maximum of 3 artificial data points across all 34 human comparisons (data not shown).

To determine biological relevance of the 1525 genes identified, pathway enrichment analysis was performed using g:Profiler ^35^ with KEGG, WikiPathways, Reactome and human phenotype ontology databases (**Fig. 2**). We selected the first 10 enriched terms based on the number of intersections between the genes in the input list and those in the enriched pathway from the respective databases. The enriched terms were related to neurological disorders and metabolic pathways whose association with PD was previously identified. Specifically, we found terms such as ‘Alzheimer’s disease’, ‘Parkinson’s disease’, ‘Huntington’s disease’, ‘Pyruvate metabolism and citric acid cycle’, ‘Regulation of lipid metabolism’, and ‘Abnormality of the nervous system’ among others. Although not captured in the top-ten list, other PD related features such as ‘dystonia’, ‘abnormality of movement’ and ‘motor delay’ were also significantly enriched. The 1525 gene panel achieved a 60% prediction accuracy when tested on an independent blood-derived transcriptome data ^36^ using random forest classifier with 10-fold cross-validation (**data not shown**); which is a relatively good performance when compared with other gene expression-based classification models in the literature ^36,37^. This further suggests that the selected gene panel is biologically appropriate for studying PD.

**Fig. 1:**
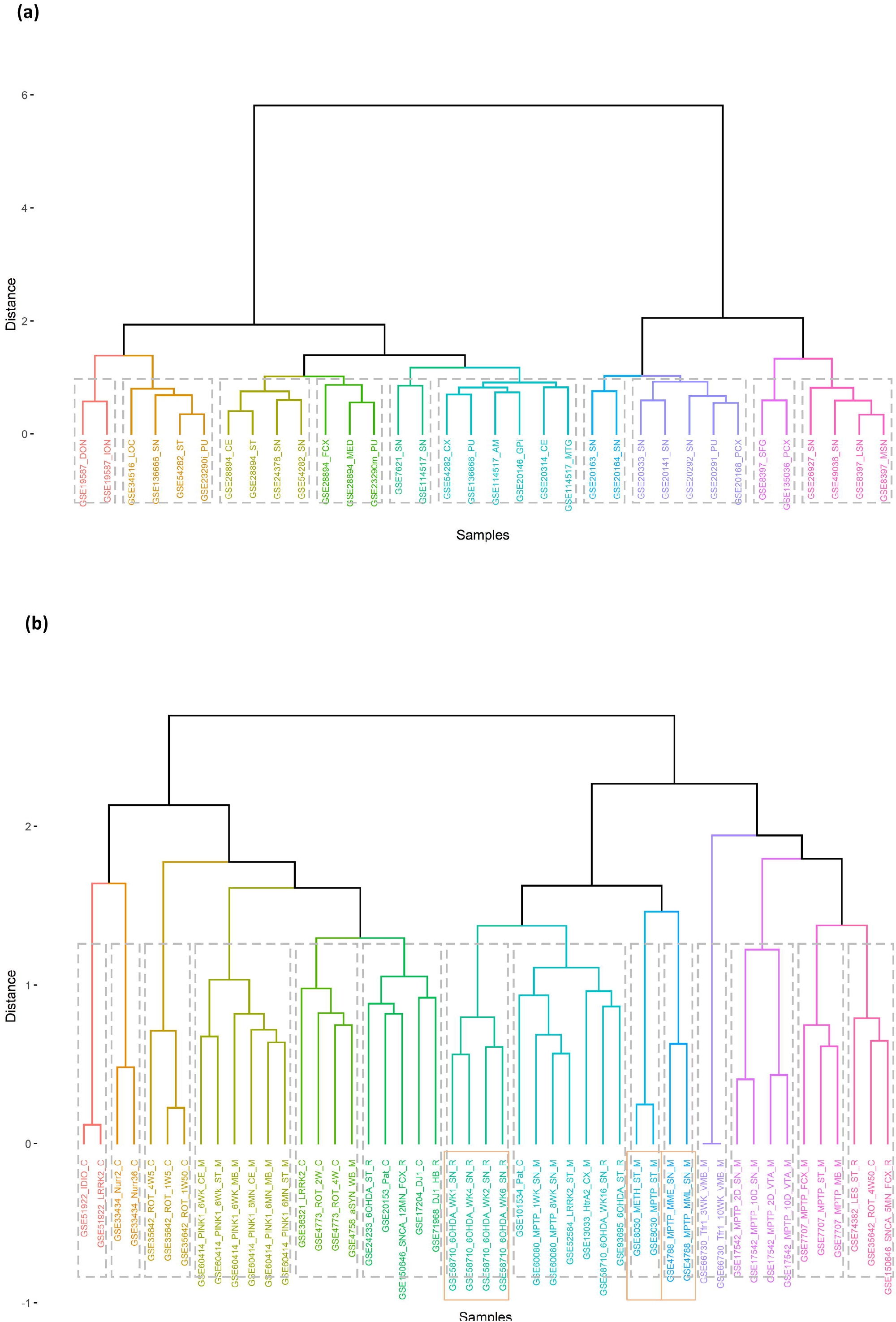
A schematic illustration of the main steps followed in the study. All the data used at each step are publicly available. GEO – gene expression omnibus, gs – gene score, FC – gene fold change, P.Value – gene p-value (from empirical Bayes moderated t-test).

**Fig. 2:**
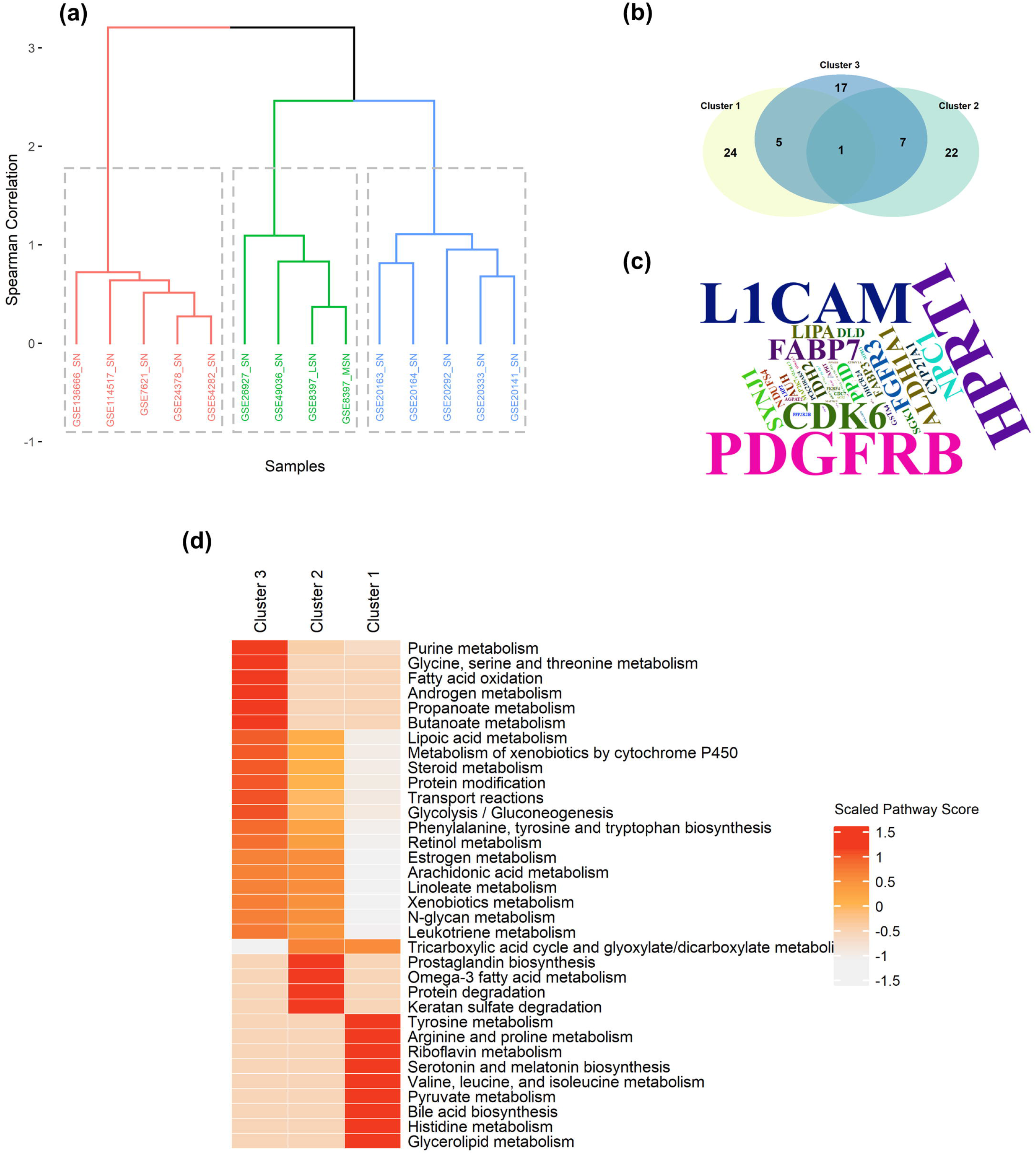
Enriched pathways in the list of 1525 metabolic genes. The first 10 enriched pathways from KEGG (Kyoto Encyclopedia of Genes and Genomes), REAC (Reactome), WP (WikiPathways) and HP (Human Phenotype) databases are shown. The size of interaction (x-axis) was z-transformed using mean and standard deviation. FDR: False discovery rate.

### Stratification of PD using transcriptome data-derived metabolic gene scores

Using the gene score data, we first sought to identify brain regions that were metabolically similar during PD. Spearman correlation clustering using the selected gene panel showed two main clusters. Within these clusters, one consisted mainly of SN and cortical derived tissue cohort datasets while the other was composed of multiple tissue type datasets, derived mainly from the hindbrain, brainstem, and other non-SN midbrain neurons (**Fig. 3a**). The same gene panel was able to distinctly group some experimental model comparisons in a brain region-specific manner, especially for striatum and substantia nigra datasets (**Fig. 3b**). However, most of these model-derived datasets appeared to have laboratory-specific effects, implying potential inherent noise within each study (**Fig. 3b**). Also, the animal models comprised both genetically and chemically induced parkinsonism with different disease durations, which might have led to different degrees of molecular effects on the level of gene expression or targeted molecular networks and might have contributed to these variations. Some cell-line models grouped with animal models (**Fig. 3b**). Interestingly, human cell-line models also resulted in non-uniform grouping of human samples (**data not shown**) when analysed together and, thus, were analysed separately. We also checked whether the experimental models would cluster with human tissues in a brain-region specific manner but found separate clusters for the two using the same feature gene panel.

**Fig. 3:**
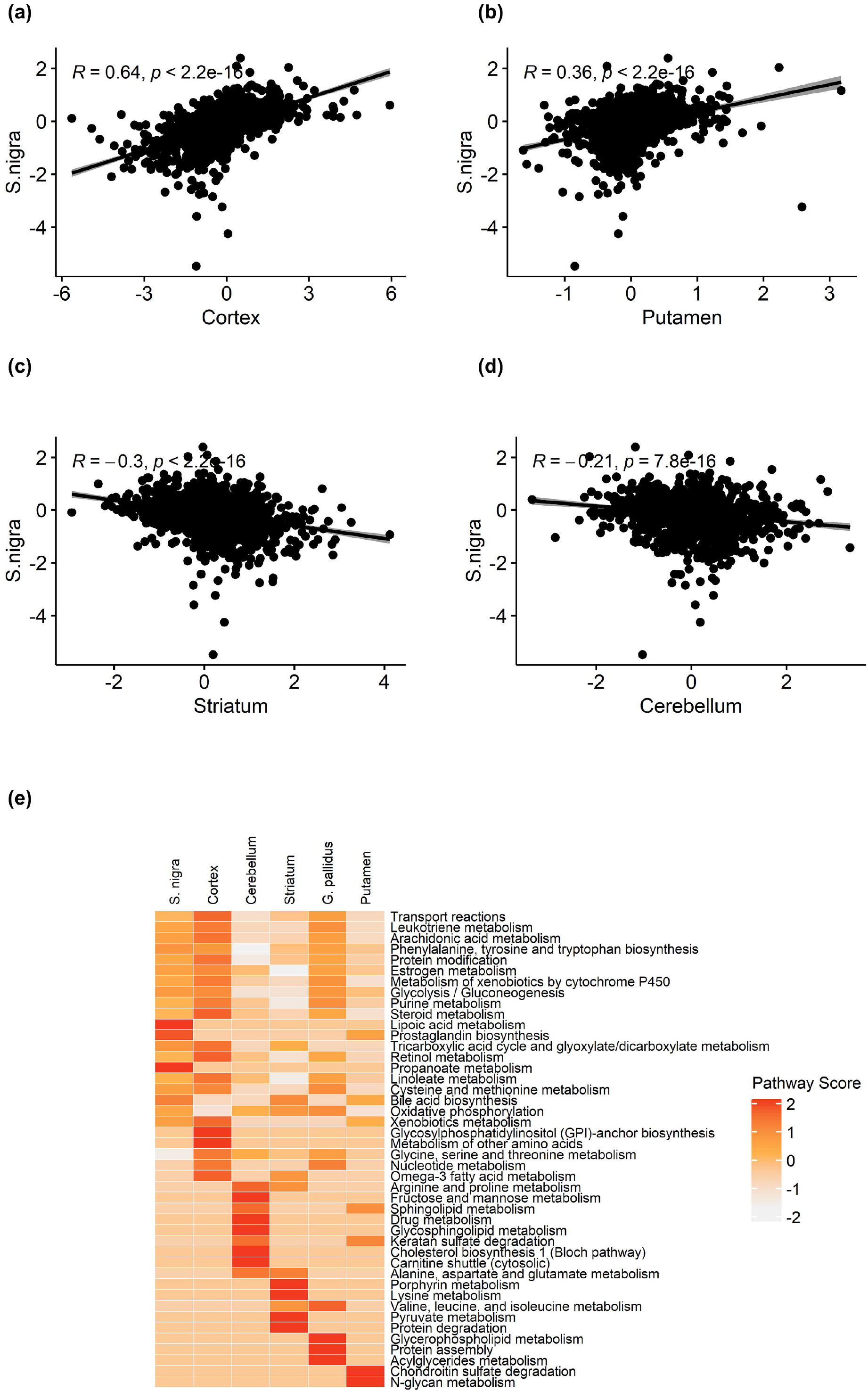
Hierarchical clustering of brain tissues. Spearman’s correlation on metabolic gene scores in **(a)** human brain tissues with most SN and cortical tissues in one major group and **(b)** experimental models (cell lines, mouse, and rat) showing metabolically closely related tissues. Closely related tissues/groups are indicated by shorter dendrogram heights. The correlation in experimental models was computed using 729 genes obtained after filtering for 95% genes with expression values and were a subset of the 1525 genes obtained from the human tissues. Pat – patient derived in vitro models. Chemical models: ROT-rotenone, 6OHDA – 6-hydroxydopamine, MPTP –1-mthyl-4-phenyl-1,2,3,6-tetrahydropyridine, METH – methamphetamine; Genetic models: IDIO – idiopathic (no known genetic association), LRRK2 – leucine rich repeat kinase 2, Nurr – nuclear receptor, HtrA2 – mouse orthologue of human HTRA2 gene, aSYN & SNCA – alpha synuclein, Tfr1 – transferrin receptor 1, DJ-1 – DJ1 protein deficient.

In summary, these observations suggest that in terms of metabolic gene scores: (i) SN is more highly correlated with CX than the other regions in the basal ganglia in the human brain (**Fig. 3a**), (ii) while less represented across the captured human datasets in this study, striatum, globus pallidus and putamen in the BG are highly correlated, but, are somewhat generally less correlated with SN (**Fig. 3a)** (iii) gene score captures and possibly conserves gene specialization in different brain tissues and might be a better approach in analysing effects of a biological perturbation across multiple tissues (**Figs. 3a and b**), and (iv) the patterns of gene scores in PD experimental models might be different from those in corresponding brain region obtained from post-mortem human brain transcriptome data.

### Comparison of Substantia nigra PD patient cohorts and associated metabolic alterations

SN was the most represented brain region from our GEO/ArrayExpress database search, with 14 comparisons from 13 separate transcriptome cohort studies (**Supplement 2** (**Table S1**)). Moreover, it is the most affected brain region displaying the highest neuronal degeneration during PD ^12^. Therefore, we investigated for the existence of cohorts with similar metabolic alterations in PD as this might inform proper patient classification based on their location for effective therapeutic interventions and group specific metabolic biomarker identification. Unsupervised hierarchical clustering using non-parametric Spearman’s correlation detected three main groups when 1103 SN specific genes were used (**Fig. 4a**).

**Fig. 4:**
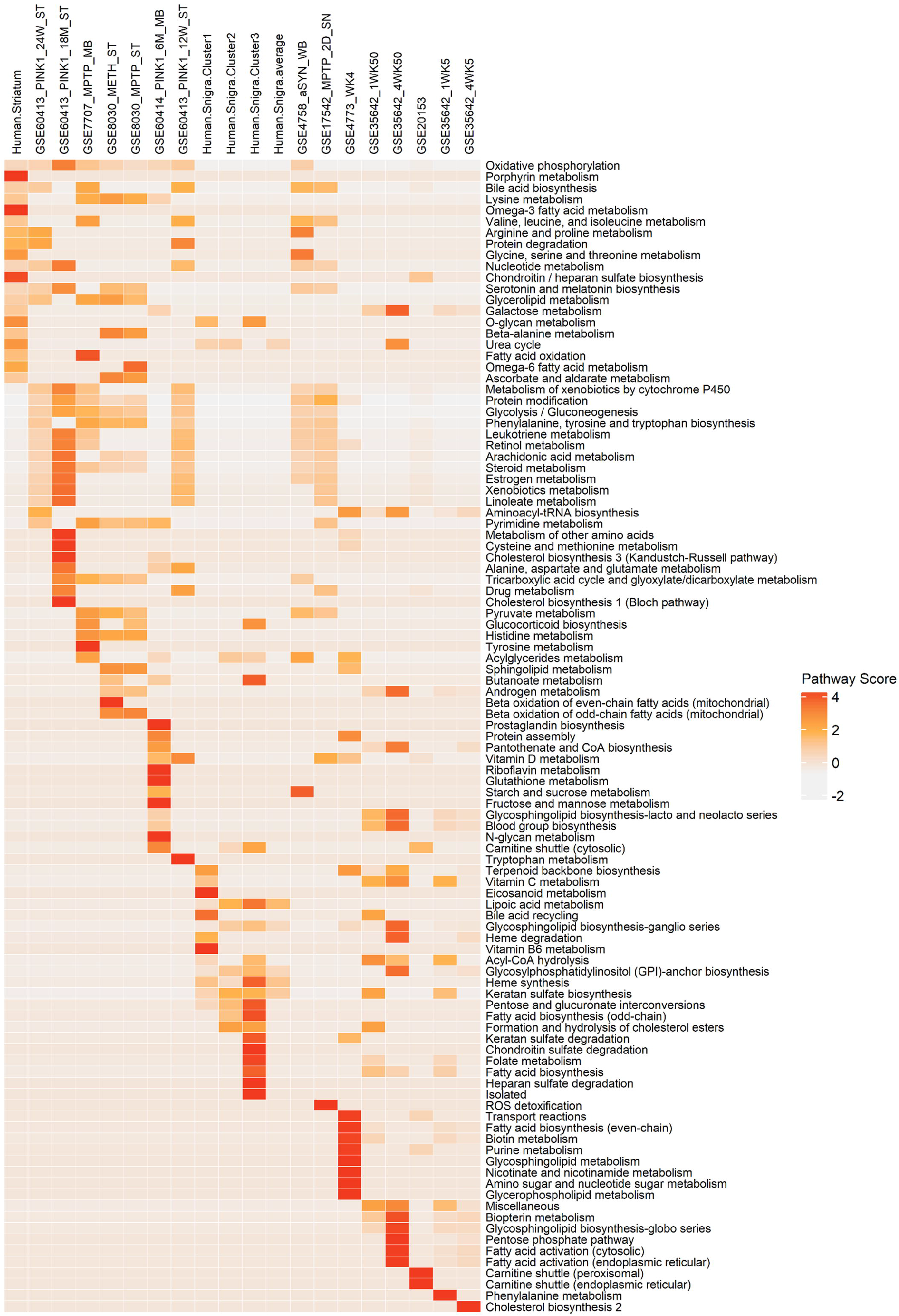
Characterization of metabolic clusters in human Substantia nigra. a) Hierarchical clustering using Spearman’s correlation as the distance measure showing three main SN related groups. b) Venn diagram describing the intersection of genes pooled from the top 20 of every group. c) Wordcloud plot showing the frequency with which the prioritized genes has been mentioned with either PD or aging in PubMed abstracts. The bigger the size the higher the frequency. d) Heatmap of pathway scores showing pathway activity for the first 20 pathways ranked based on pathway scores in each of the three detected SN groups.

We next sought to determine the molecular differences and biological relevance of these clusters based on the gene scores of the 1103 SN specific genes. For each cluster, we ranked the average gene scores and prioritized the first/last 15 genes (30 genes in total per group) (**Table 2**). We found each cluster to contain some unique genes as well as other overlapping genes (**Fig. 3a**). Next, we performed literature search to check if there were existing associations between these genes and PD/aging (aging is one of the risk factors of PD) ^6^. Out of 76 unique genes, we found 64 (84%) genes to have associations with either the aging process or PD or both, and 27/64 (42%) genes had been associated with only the aging process from PubMed literature reports. Within the aging-only associated genes, 6/27 genes were directly associated with lipid while 2/27 mitochondrial metabolism respectively – metabolic processes believed to be perturbed by PD pathology ^1,4,5,10,11,38^. **Fig. 4c** shows a wordcloud plot showing the frequency with which some of the genes were associated with PD in PubMed publications. RNF130, B3GALT2, GUCY1A2, CYB5R1, POLR2L, ALD3B1, SLC6A12, PTDSS1, PIP5K1B, MAN2A1, B4GALT6 and CDK14 (12) genes had no explicit association with PD. A summary of these genes pertaining to their names and PubMed article IDs of the studies they appear is provided as **Supplement 2 (Table S2)**. When the clusters were compared (**Fig. 4b**), ALDH1A1 was found to be common to all the groups. ALDH1A1 gene belongs to a family of detoxification enzymes and is involved in the metabolism of catecholamines, including dopamine and norepinephrine, and detoxifies aldehydes in the SN ^39,40^. A meta-analysis with 6 separate transcriptome datasets suggested this gene to be downregulated in the SN ^41^. Indeed, we observed that it had a negative gene score across all the SN groups, with a relatively high score magnitude in the third group, suggesting that its effect might be more pronounced in some patient cohorts. Unfortunately, it was not possible to establish whether enrichment of this gene was correlated with previous levo-dopa use across all groups. There were no unique genes common to groups 1 and 2 in this list. However, between clusters 1 and 3, we found 5 genes (RNF130, CA2, ABCA8, CLK1 and FABP7) while clusters 2 and 3 had 7 genes (SLCO4A1, PTDSS1, SYNJ1, ACOT7, PIP5K1B, PRKAR2B, and GBE1) in common. These genes were mainly associated with cellular protein clearance/degradation, transport, RNA metabolism, neuron cell development, learning and catalytic activities. We further checked the overlap between the prioritized genes with those recently proposed from a meta-analysis of differentially expressed genes in the human SN ^41^ vis-à-vis their association with Mendelian diseases as contained in the OMIM ^42^ disease database. PLOD3 and LRP2 were in common with up-regulated gene list, while AUH, ATP6V1A, PPP2R2B, SYNJ1, DLD, PTDSS1, GBE1, L1CAM, and HPRT1 were down-regulated in PD versus HC comparisons, respectively in the study, suggesting that the prioritized gene lists from the metabolically similar study cohorts play important metabolic roles in the SN during PD and defective forms might be inherited/deregulated through mutations or environmental factors increasing the odds of developing PD. It will be important to systematically evaluate the impact epigenetics might play on their dysregulation to fully understand how they come about in PD.

**Table 1:** Number of transcriptomic datasets, comparisons and samples used in this study

|  | Human<br>Subjects | in vivo<br>mouse<br>models | in vivo rat<br>models | in vitro<br>human<br>models | **TOTAL** |
| --- | --- | --- | --- | --- | --- |
| Number of datasets | 21 | 11 | 6 | 9 | **47** |
| Number of comparisons | 34 | 34 | 10 | 14 | **92** |
| Number of control / PD<br>samples | 285 / 334 | 93 / 106 | 46 / 46 | 51 / 67 | **1028** |

**Table 2:** Top 30 highly ranked genes based on their absolute gene scores from each Substantia nigra metabolic gene expression group.

| Cluster 1 |  | Cluster 2 |  | Cluster 3 |  |
| --- | --- | --- | --- | --- | --- |
| Gene Symbol | Gene Score | Gene Symbol | Gene Score | Gene Symbol | Gene Score |
| SLC5A3 | 2.817 | SLCO4A1 | 2.289 | ABCA8 | 6.850 |
| RNF130 | 2.154 | BCAT2 | 2.200 | SLCO4A1 | 6.455 |
| FKBP4 | 2.135 | MAP2K7 | 2.046 | RNF130 | 6.433 |
| PLOD3 | 1.994 | MAP3K11 | 1.915 | CTDSP2 | 5.896 |
| CA2 | 1.697 | SHMT2 | 1.850 | CYP27A1 | 5.734 |
| PPID | 1.530 | TCIRG1 | 1.812 | NPC1 | 5.362 |
| CDK6 | 1.474 | ISYNA1 | 1.702 | PIP4K2A | 5.266 |
| KIT | 1.440 | ALDH3B1 | 1.658 | SGK1 | 5.249 |
| B3GALT2 | 1.398 | PTPRO | 1.622 | CA2 | 5.139 |
| ATM | 1.390 | PLCB3 | 1.557 | CLK1 | 5.009 |
| CDC7 | 1.384 | SLC6A12 | 1.548 | PPM1B | 4.949 |
| ABCA8 | 1.377 | IRAK4 | 1.449 | MAN2A1 | 4.698 |
| GUCY1A2 | 1.298 | AGPAT2 | 1.416 | LRP2 | 4.664 |
| DHCR24 | 1.261 | PDGFRB | 1.404 | PLCL1 | 4.529 |
| CLK1 | 1.252 | POLR1B | 1.394 | LIPA | 4.462 |
| DECR1 | -1.246 | AUH | -3.273 | HPRT1 | -5.427 |
| IDH2 | -1.247 | PTDSS1 | -3.285 | B4GALT6 | -5.491 |
| CYB5R1 | -1.258 | NDUFS4 | -3.336 | PTDSS1 | -5.595 |
| RBP1 | -1.270 | ATP6V1A | -3.378 | PTPRN | -5.739 |
| FABP3 | -1.314 | SYNJ1 | -3.440 | FABP7 | -6.799 |
| LIMK2 | -1.317 | UQCRC2 | -3.465 | ACOT7 | -6.856 |
| GGT5 | -1.318 | GSTA4 | -3.551 | ACHE | -6.954 |
| ACP2 | -1.358 | DLD | -3.621 | HMGCS1 | -6.976 |
| POLR2L | -1.376 | PPP2R2B | -3.710 | SYNJ1 | -7.117 |
| PLCD1 | -1.401 | MDH1 | -3.852 | L1CAM | -7.985 |
| FGFR3 | -1.405 | ACOT7 | -3.957 | CDK14 | -10.501 |
| FABP7 | -1.464 | PIP5K1B | -4.215 | PRKAR2B | -11.480 |
| APRT | -1.718 | PRKAR2B | -4.390 | PIP5K1B | -13.338 |
| PCK2 | -1.724 | GBE1 | -6.593 | ALDH1A1 | -14.082 |
| ALDH1A1 | -3.485 | ALDH1A1 | -9.621 | GBE1 | -14.367 |

A further interrogation of these clusters through metabolic pathway score analysis was performed to functionally characterise the differential enrichment of metabolic pathways specific to each cluster. For each cluster, the average p-value and fold change of each gene across all comparisons was computed and used to calculate pathway scores. Top 20 pathways from each cluster based on maximum pathway scores across all comparisons in each cluster were used. In **Fig. 4d**, cluster 1 showed stark pathway score differences from clusters 2 and 3. This difference was mainly from high pathway scores for amino acids (tyrosine, histidine, arginine, proline, valine, leucine and isoleucine), pyruvate, bile acid, glycerolipid, riboflavin and serotonin and melatonin metabolisms. Clusters 2 and 3 shared most of the active pathways such as leukotriene, lipoic acid, steroid, arachidonic acid, linoleate, oestrogen, retinol, xenobiotic, N-glycan, protein modification, transport, glycolysis/gluconeogenesis, phenylalanine, tyrosine and tryptophan metabolisms. Additionally, these clusters were differentially active for prostaglandin, omega-3 fatty acid, protein degradation and keratan sulphate (cluster 2) and fatty acid oxidation, propanoate, butanoate, purine, glycine, serine, and threonine (cluster 3) metabolisms. The lipid metabolism altered in these clusters relate to energy generation (fatty acid oxidation, propanoate and butanoate) and immune system function (prostaglandin and omega-3 fatty acid). Some of the metabolic pathways found in these clusters have been extensively studied for their association with PD and are believed to be important biomarkers of the disease. For instance, perturbations in energy metabolism through mitochondrial dysfunction ^4^ and dysregulated lipid metabolism ^10,43^ have been experimentally validated in PD. Other metabolites such as Serotonin, melatonin and 2-oxobutanoate were amongst the metabolic changes recently proposed to be PD biomarkers in mouse models ^44^. In addition, a recent study on Alzheimer’s disease (AD) demonstrated strong perturbations in bile acid metabolic pathways ^45^. Given the close aetiologic relationships between these diseases, our finding on the enrichment of this pathway might suggest that this pathway is not unique to AD and this is supported by other findings^1,5,10,11,46^.

Altogether, these findings show multiple cluster-specific metabolic perturbations in the SN. The clusters indicate subsets with perturbations in the (i) neurotransmitter, energy, and complex lipid (bile acids and glycerolipids) metabolisms, (ii) metabolism of inflammatory response lipids and (iii) lipid breakdown for energy metabolism.

### Regional brain metabolic correlation within the BG-CX-CE axis in PD

Initiation and execution of motor and non-motor functions in the brain rely on a well-defined regional brain coordination circuitry that is frequently disrupted in PD and contributes to clinical symptoms of the disease. We checked how PD alters metabolic genes within the BG-CX-CE molecular circuitry ^13^ by performing gene correlation analysis on average gene scores from each brain region using Spearman correlation. This axis is embedded with substantial neuronal projections and, additionally, are spatially close in the human brain. In each analysis, we compared the average gene scores between SN tissues and cerebral cortex, putamen, striatum and cerebellum – tissues belonging to the BG-CX-CE axis. The choice of SN as the central point of our analysis was informed by the fact that it undergoes the most severe degeneration and thus highly significant in PD ^13^. Also, other studies have suggested that the resultant degeneration of these neurons might impact the biological functions of dependent neurons for executing coordinated tasks through reduced or absent neuronal neurotransmitter signals ^2,13^.

Significantly strong positive correlations were found between SN and cerebral cortex (*ρ* = 0.64), putamen (*ρ* = 0.36). In contrast, negative correlations were found with striatum (*ρ* = −0.3), and cerebellum (*ρ* = −0.2) (all Student’s t-test *p* < 0.05) (**Figs. 5a, b, c and d**). The findings of a significant negative correlation between SN and striatum (**Fig. 5c**) and a positive correlation with putamen (**Fig. 5b**), which are tissues within the BG and are known to influence different PD phenotypes ^13^, was intriguing and might suggest that neuronal degeneration during PD extends to the putamen but less to striatum. This finding augurs well with a recent finding from a mouse model suggesting the independence of these regions ^47^. A strong local interconnectivity between SN and cerebral cortical tissues has recently been shown ^2,48^ and the findings here suggest that such interconnectivity is also present at the differential expression of metabolic genes level. On the other hand, the correlation with cerebellum, a member of the BG-CX-CE circuitry, was found to be negative (**Fig. 5 d**) suggesting that the impact of the pathology on SN might be having a negative influence on these tissues.

**Fig. 5:**
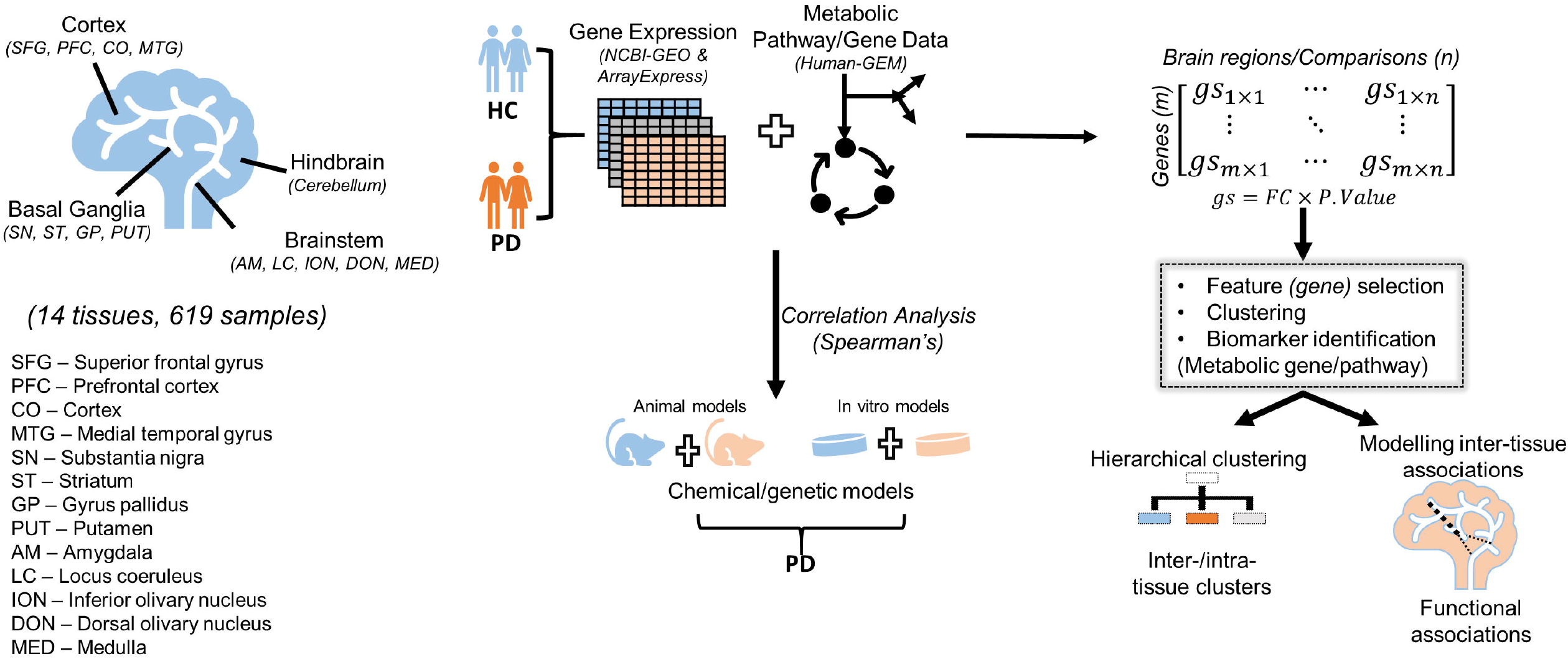
Metabolic gene correlation in the BG-CX-CE axis during Parkinson’s disease. a-e) Scatter plots showing the linear regression and correlation between Substantia nigra and **(a)** cerebral cortex, **(b)** putamen, **(c)** striatum, **(d)** cerebellum respectively (p<0.05). **(e)** Heatmap of the top 20 high scoring pathways from each brain tissue illustrating the differences and similarities in pathway activity across the brain tissues. The points correspond to each gene whose scores were compared.

Additionally, we analysed the correlation of metabolic gene scores between SN and other tissues beyond the BG-CX-CE axis that were included in our study. Tissues in the brainstem (ION, DON, ME) are known to be first affected by α-Syn pathology believed to arrive from the gut in PD through vagus nerve endings (brain-gut axis). They also lie immediately below the BG. We found low correlations with ME. ME contains both ION and DON specific cell types. When we analysed these cell types separately, weak but significant positive correlations were found with the DON (*ρ* = 0.13) and not ION suggesting specificity of metabolic alterations might be down to the neuronal cell type level in the brain regions first affected in PD.

To gain more insight and determine the biological significance of the correlation analysis, pathway score analysis was performed, and the top 20 pathways based on average pathway scores for each brain region was used to identify the functional metabolic differences/overlaps. From our analysis, the high correlation between SN and CX, for instance, might be due to a higher relative metabolic pathway scores for protein modifications, transport reactions, nucleotide metabolism, lipids (steroid, oestrogen, leukotriene, arachidonic and linoleate) and glycolysis/gluconeogenesis metabolism, which had relatively similar activities in these tissues. The negative correlation between metabolic gene correlation in SN versus striatum and CE might be due to the opposite pathway scores on amino acid metabolism (arginine and proline and lysine), lipids (sphingolipid and glycosphingolipids and cholesterol biosynthesis), pyruvate, porphyrin, keratan sulfate, carnitine shuttle and protein degradation metabolism.

Thus, our findings suggest that the metabolic alterations in the SN correlates more with the corresponding alteration patterns in the cerebral cortex tissues than members of the BG (putamen, globus pallidus and striatum) and CE within the BG-CX-CE functional molecular circuitry. These observations also correlate with the clustering analysis findings (**Fig. 3**) where most SN and CX comparisons clustered closer as compared to other regions included in this analysis and suggests that the closeness between these regions might be due to similar metabolic alteration patterns resulting from α-Syn pathology. However, future studies using longitudinal multi-omics data might shed more light on these findings as most of the data we used in this study were from separate studies and environmental factors contributing to PD might have different effects on the tissues studied.

### PD disease model selection based on metabolic gene scores

Due to the delicate nature of the human brain inhibiting invasive experimental sample collection, experimental models are of upmost importance and using the right model to simulate an intended biological process is equally important. Since we had observed substantial metabolic alterations in the various PD cohorts, we checked whether available experimental PD models (human-derived cell lines, rat, and mouse), both chemically and genetically induced, would be suitable for systematic modelling of metabolism during PD.

We performed correlation analysis using average gene scores between human brain tissues and the corresponding tissue in the experimental models. Aside from SN and striatum tissues, there were insignificant correlations between human and corresponding experimental model gene expression. For SN models, correlations were performed separately with each of the 3 previously found cohorts (**Fig. 3a and Supplement 1 (Figs. S2a and b)**) to allow us to identify human SN clusters that were metabolically highly similar to the experimental models. The other brain regions, notably striatum and PCX, were directly compared as no clusters had been defined. For both cell lines and mouse experimental models, low correlations (<0.5) were found mostly with mouse models. Interestingly, the amount of correlation differed between the different clusters of SN. For instance, while 1 week rotenone treatment (5nM and 50nM) showed negative and positive correlations with the first and second SN cohorts, 4-week treatment had an opposite trend on gene scores in the GSE35642 cell line dataset. GSE4773 and GSE20153 were more correlated with the second SN cohort. Indeed, most metabolic pathways active in the SN are inactive in the cell line experimental models, except for a few pathways such as lipoic acid, bile acid recycling, carnitine shuttle, acyl-CoA hydrolysis, urea cycle, keratan biosynthesis metabolism, among others (**Fig. 6**). Most of these pathways are related to the lipid metabolic pathway suggesting that the complete repertoire of altered metabolic pathways in PD as found in the human tissues (**Fig. 4d** and **Fig. 5e**) is partially covered.

**Fig. 6:**
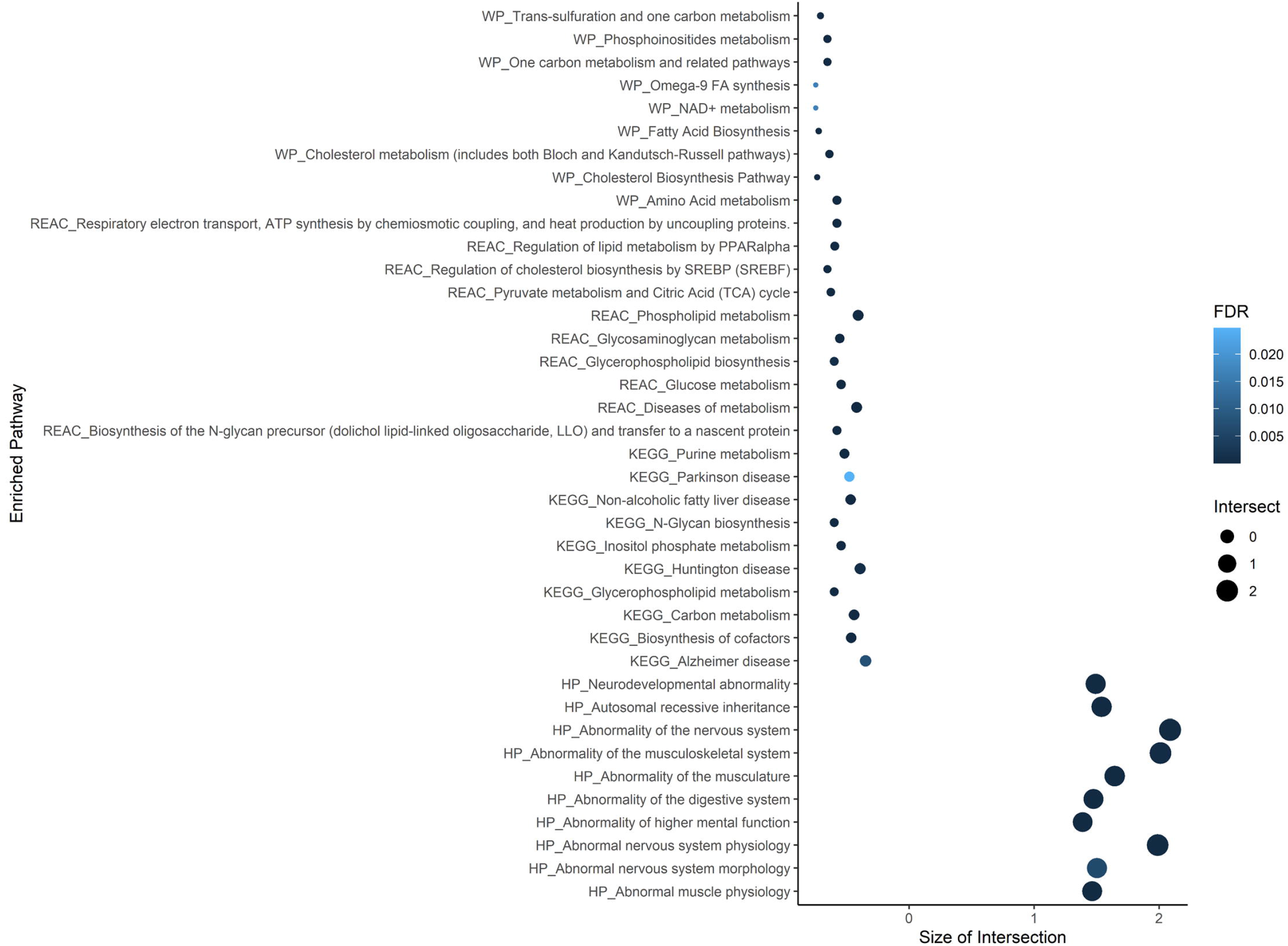
Comparison of metabolic pathway scores between human PD tissues and experimental models. Heatmap of metabolic pathway scores showing metabolic functions whose activities might be shared with human tissues as well as their metabolic differences. The depicted models are those that had an appreciably good absolute correlation score (ρ≥0.1).

Except for GSE7707 dataset, all striatum mouse models were found to have a negative correlation with the corresponding human tissue, regardless of the modelling technique used (i.e., chemical, or genetic) (**Table 3**). Accordingly, **Fig. 6** shows that most of the pathways with increased activity in these experimental models are indeed inactive in the corresponding human striatum tissue.

**Table 3:** Experimental models with significant correlations with modelled human tissue. All reported results had absolute correlation coefficient (ρ) ≥ 0.1 and Student’s t-test p-value < 0.05.

| Tissue | Model system | Experimental Model Dataset | Spearman's Correlation | p-value |
| --- | --- | --- | --- | --- |
| SN | Cell line | GSE4773_ROT_WK4 | -0.14 (cluster2) | <0.05 |
|  |  | GSE20153_Pat | 0.25 (cluster2) |  |
|  |  | GSE35642_ROT_1WK5 | -0.18/0.17 (cluster1/2) |  |
|  |  | GSE35642_ROT_1WK50 | -0.2/0.28 (cluster1/2) |  |
|  |  | GSE35642_ROT_4WK5 | 0.17/-0.22 (cluster1/2) |  |
|  |  | GSE35642_ROT_4WK50 | 0.2/-0.22 (cluster1/2) |  |
|  | Mouse | GSE4758_WB | -0.13 | <0.05 |
|  |  | GSE17542_2D_SN | 0.1 |  |
| ST | Mouse | GSE7707_MPTP_MB | 0.12 | <0.05 |
|  |  | GSE8030_MPTP_MB | -0.14 |  |
|  |  | GSE8030_METH_MB | -0.21 |  |
|  |  | GSE60414_6M_MB | -0.13 |  |
|  |  | GSE60413_12W_ST | -0.21 |  |
|  |  | GSE60413_24W_ST | -0.12 |  |
|  |  | GSE60413_18M_ST | -0.11 |  |

SN: Substantia nigra, ST: Striatum, MPTP: 1-methyl-4-phenyl-1,2,3,6-tetrahydropyridine, METH: Methamphetamine, ROT: Rotenone, WB: Whole brain, W: Week, Pat: Patient, D: Day, MB: Mid-brain.

In summary, while modelling human metabolic alterations in PD, only a few available experimental models might recapitulate the molecular metabolic landscape in humans. Even those with significant positive correlations were merely moderate. This is an interesting finding since current studies exclusively rely on these experimental models to characterise the biology of PD and prioritize potential therapeutic interventions. Thus, there is an urgent need to fully determine the exact molecular differences between human and experimental models of PD which would inform better model creation for future studies. Nevertheless, given the underlying disease molecular complexity in PD, the roles played by epigenetics as relates to the development of most cases of idiopathic PD might be difficult to reproduce in these models under experimental conditions but might greatly improve the accuracy of the altered molecular processes in these models with potential benefits for PD drug research and potential disease biomarker evaluation.

## Discussion

The analysis pipeline adopted in this work demonstrated that aggregation of transcriptome data using gene scores approach and filtering for genes with high variability result in the discovery of disease relevant molecular features. In total, we analyzed 92 comparisons from 47 transcriptome datasets, covering 1028 samples in total. We analysed the expression of metabolic genes from post-mortem human brain during PD to identify (i) metabolically related brain regions, (ii) SN clusters from PD patient cohorts displaying similar metabolic alterations and (iii) differences and similarities in metabolic alterations associated with BG-CX-CE circuitry in PD. Finally, we used the identified clusters to identify experimental models (cell line, mouse, and rat) and characterised the metabolic alterations they would model. We found the existence of metabolic alteration heterogeneity in PD brain regions across study cohorts, particularly for the SN with three main clusters.

In our simulations, using either gene fold change or p-value alone in clustering analysis did not give meaningful clusters for the same brain region from different study cohorts. However, by combining the two and computing a gene score it was possible to identify disease relevant genes (**Fig. 2**) and find clusters of similar brain regions (**Fig. 3a**). The gene list captured biological processes related to energy, neurotransmitter, protein processing and inflammation metabolism, which are relevant in PD ^4^, thereby suggesting the importance of the two measures in delineating brain tissues from their gene expression profile. Using gene scores, we observed high correlations in the expression patterns of metabolic genes between the SN and cortical tissues as compared to other tissues in the basal ganglia region (**Fig. 3a** and **Figs. 5a, b**, and **c**). The SN-CX high correlation is consistent with previous suggestions on the close coordination of these tissues, which is a target of PD pathology ^13^. We adopted a similar strategy to identify three clusters of SN from multiple study cohorts (**Fig. 4a**) and observed a low gene overlap between the clusters (**Fig. 4b**). Subsequently, we identified experimental models with significant correlations with human brain tissues based on their gene scores. Thus, gene scores approach yielded consistent results and might be a good approach in systematic analysis relying on gene expression data from different high-throughput data collection platforms or separate studies.

By computing metabolic pathway scores, we were able to biologically characterize the observed inter-region brain correlations and clusters derived in each analysis. For instance, we observed perturbations in lipid metabolism and glycolysis (**Fig. 5e**) to correlate with the observed high molecular similarities between SN and CX. Indeed, previous findings have found these alterations in PD patients ^1,5,49,50^. Similarly, we established the metabolic perturbations contributing to the observed differences between SN and (i) striatum (in the basal ganglia) and (ii) CE (**Fig. 5c** and **d**), for which we had found negative correlations. Interestingly, a recent systematic analysis had suggested that while α-Syn as Lewy bodies accumulates in several brain regions during PD progression ^12^ its impact on gene expression is varied, with some neurons being more vulnerable and undergo degeneration ^51^. Altogether, this characterisation highlighted a region-specific metabolic dysregulation during PD and suggested that different metabolic biomarkers might be important for assessing the severity of PD at a brain-wide scale. Moreover, metabolic biomarker diversity can distinguish different patient cohorts based on disease perturbations on the SN as we observed (**Fig. 4d**). Unfortunately, only a subset of the identified metabolic alterations, such as amino acid, neurotransmitter and energy metabolism, is captured by genetic or chemical (or a combination of both) modification of experimental models (**Fig. 6**). Altered metabolites related to these processes were recently identified in a metabolic profiling of PD-induced mouse brain ^52^.

This study has following limitations that need to be addressed by subsequent studies: (i) we did not control for sample covariates such as gender, age, Braak stage ^12^ etc., which are known to directly influence gene expression patterns, (ii) the samples were collected from patients from different geographical locations and environmental factors contributing to PD might differ, and (iii) the molecular evidence presented are based on predictions from one omic layer, thus longitudinal study with a multi-omic approach might provide cohort-specific reproducible molecular alterations in a PD, which is a major challenge facing PD research currently ^50^.

In conclusion, the findings from this study have potential implications on PD drug research and personalized medicine as they enforce that (i) PD is not a single disease, from a metabolic perspective, but consists of multiple clusters with each having a unique biomarker repertoire that should be taken into consideration while devising a therapeutic strategy, (ii) SN and CX brain regions are the most metabolically interdependent regions during PD meaning that this interconnectivity present a potential avenue for new drug target and biomarker identification, and (iii) only a handful of currently available PD experimental models can mirror some aspects of metabolic alterations during PD and these might be biased to a few patient subgroups/cohorts. However, whether the observed low correlation between experimental models and corresponding human tissues is a function of disease stage or due to biological differences between the organisms still need further investigation.

## Methods

### Transcriptome data acquisition and pre-processing

Raw transcriptome datasets from studies on brain tissues of human Parkinson’s disease (PD) patients and from genetically or chemically induced PD cell line and animal (rat and mouse) models were downloaded from Gene Expression Omnibus ^19^ (https://www.ncbi.nlm.nih.gov/geo/) and Array Express ^20^ (https://www.ebi.ac.uk/arrayexpress/) databases using GEOquery ^21^ and ArrayExpress ^22^ R packages respectively. To ensure statistical soundness of each comparison, we only chose studies that had at least 3 samples in both disease and control groups. Transcriptome dataset accession numbers and other related information are given in **Supplement 2 (Table S1)**. Oligo and lumi packages in R were used to process raw Affymetrix and Illumina microarray datasets respectively. Robust Multi-chip average (RMA) method in the Oligo package was used for background correction, normalization, and summarization of probe arrays. Quantile normalization method was used in all cases for the normalization of raw data. Raw count data were normalized by the TMM method in edgeR package for RNA-seq derived datasets. Here, genes with less than 1 count absolute expression across all samples were removed from further analysis as they were assumed to be less informative. Some datasets included multiple brain regions or multiple PD models, yielding 92 control-PD comparisons across 47 datasets. Of these comparisons, 34, 14, 34 and 10 were from post-mortem human, human cell lines, mouse, and rat datasets respectively. The number of control and PD samples in each category is provided in **Table 1**. For microarray data, gene Entrez IDs with maximum detected signal were selected in the case of multiple platform probe IDs per gene.

Next, the list of human metabolic genes was obtained from a recent reconstruction of human genome-scale metabolic model (Human-GEM v1.3.0) ^23^. Mouse and rat orthologs of the human metabolic genes were extracted using biomaRt R package ^24^ after converting human gene Ensembl IDs in Human-GEM to their corresponding human gene Entrez IDs. This Human-GEM model contains 3628 Ensembl metabolic gene IDs ^23^ with 3634 corresponding Entrez IDs, and the corresponding mouse and rat metabolic gene Entrez IDs identified by the ortholog analysis are 3865 and 3899 respectively. All subsequent analyses were performed using gene Entrez IDs and corresponding HGNC gene symbols. Metabolic gene expression profiles from the transcriptome datasets were extracted using these gene lists.

Principal component analysis (PCA) ^25^ was then used, with the first two principal components, to visually detect and remove outlier samples from disease versus healthy controls comparison in both human and model samples. Gene fold change and empirical Bayes p-values between disease and control groups were calculated using limma ^26^ package in R. Gene fold changes for down-regulated genes were represented as -1 divided by the corresponding gene fold change to make them on the same scale as up-regulated genes. The sign was introduced to distinguish up-from down-regulated genes. Gene fold change values and their corresponding p-values from all comparisons were merged into separate data matrices.

### Imputation of genes with missing expression values in a subset of studies

To avoid losing important genes that might not have been captured by a study, we applied random forest regression missing value imputation using missForest R package ^27^ for the missing data points in the combined fold-change data matrix. The algorithm was set to use 15 trees and 200 iterations to impute missing values for the genes with at least 95% real values across all comparisons, in effect discarding genes with higher than 5% missing values in the data matrix. All other parameters in the imputation algorithm were used in their default values. This led to a total of 1833 metabolism related genes across 34 comparisons from human datasets. This imputation was performed on the p-value data matrix as well. A similar imputation strategy was adopted on disease model data separately.

### Gene score and gene (feature) selection

To conserve the statistical significance and direction of metabolic gene deregulation between PD and HC, each gene was scored by combining its gene fold change and corresponding p-value into a single score for each comparison using the equation shown below [30]:

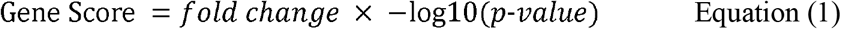

where fold change is the gene fold change, with the transformation mentioned above being valid for down-regulated genes, and p-value the corresponding empirical Bayes p-value. An unsupervised approach was further used to select genes with the highest contribution to variance in gene scores across all comparisons by using plotloadings() function in pcatools package ^28^. Here, we applied PCA to the gene scores data from human datasets, and the function selects top/bottom 20% genes that contribute to the differences along each of the principal components. PCs with at least 70% overall contribution to variance were used in each case. Given a percentage cut-off and the number of PCs to consider, plotloadings() creates a final consensus list of genes across all principal components ^28^. In this way, we avoid the gene fold change or p-value cut-off based biased approach to select variable genes across all human comparisons as they might leave out important genes that do not show expression above the specified criteria. Prior to PCA, in each case, every gene score was centred around the mean score across all comparisons and normalized with the corresponding standard deviation. The final gene list had 1525 human metabolic genes with highest variability across the selected principal components.

### Detecting metabolically correlated brain-regions in PD

Next, Spearman correlation was used to cluster 34 human comparisons based on the genes with highest variability. Ward’s minimum variance (*ward*.*D*) was used as the dis-/similarities measure between any two clusters. This was implemented using hcut() function in R. The obtained dendrogram was plotted using fviz_dend() function in factoextra ^29^ R package. This procedure was repeated separately for all the PD models (in vitro and animal) using the same gene list of 1525 genes.

### Identification of metabolic gene biomarkers for SN clusters

To prioritize genes from SN tissues, we first identified a subset of genes with high variance and unique to this tissue from the 14 SN-specific comparisons using the plotloadings() function, as explained previously, from their gene scores. Next, clustering analysis was performed with hierarchical clustering method. Then, for each identified cluster, we calculated the average gene scores across the comparisons in that cluster and ranked the genes based on whether they have high average positive or negative scores. From each cluster, the first 30 (top and bottom ranked 15) genes were selected and previous studies were checked for their relationship with PD. PUBMED abstracts were searched using the get_pubmed_ids() function with ‘[Parkinson’s] disease AND [geneX]’ or ‘[Aging] AND [geneX]’ as the keywords for previous reports on each of the prioritized genes (geneX) in PD using easyPubMed R package (available in Bioconductor database).

### Pathway score calculation for cluster functional characterization

For each of the groups detected in the clustering analysis, there exists a subset of genes with a similar expression pattern across all the comparisons in the group. To investigate the functional similarities/differences between the groups, we used metabolic pathways from the human genome-scale model, Human-GEM, (134 metabolic pathways) ^23^ to calculate pathway scores. Gene-pathway association data was extracted from Human-GEM by associating all genes involved in reactions within a metabolic pathway with that pathway. As previously described ^30^, the pathway score was defined as:

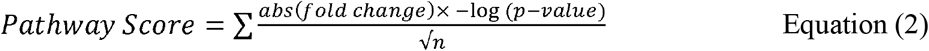

Where, *abs (fold change)* is the absolute gene fold change, *p-value* the corresponding gene p-value and *n* the normalizing factor reflecting the number of genes associated with the respective metabolic pathway. For each comparison in a group identified by clustering, we first calculated pathway scores using Equation (2), and then averaged pathway scores across the comparisons in that group to identify dysregulated pathways for each group. A graphical illustration of this procedure is provided in **Supplement 1 (Fig. S1)**.

### Analysis of metabolic correlations in the BG-CX-CE molecular circuitry in PD

BG is the main affected area during PD ^4,13^, however, disease symptoms are associated with perturbations in multiple other brain regions, which have been suggested to form a direct and indirect communication circuitry driving key motor and non-motor PD symptom progression ^13^. Furthermore, α-Syn aggregates, the main pathogenic component of LB ^4^, are known to migrate from BG to distant cells ^14^ and, therefore, disease symptoms resulting from dysfunctions in the affected sites might be indirectly explained by the eventual accumulation of the spreading pathogenic α-Syn in the target tissue ^4,12,14^.

To investigate how PD influences the BG-CX-CE circuitry at the metabolic level, we calculated the correlations between the expression changes in SN and other regions in this circuitry (CX, CE, putamen, striatum and globus pallidus), separately, using average gene scores from each region. We used Spearman correlation to determine the amount of correlation and Student’s t-test to estimate the statistical significance of the correlation coefficient. Subsequently, pathway scores were computed as defined in the previous section and visualized using a heatmap. The heatmaps were drawn using Heatmap() function in ComplexHeatmap R package ^31^.

### Disease Model Selection

Most PD animal models have been shown to be different from human PD at the gene expression level and rarely represent the underlying disease process ^17,32^. To investigate the extent to which this is true for metabolic genes, we calculated the correlation between gene scores in the human brain regions and corresponding region in experimental models. For the case of PCX and SN, we merged subgroups such as cerebral cortex (CB), LSN, MSN etc by taking the average scores, as we reasoned that those sub-classes/sub-regions would not greatly influence the correlation scores. Human orthologs of mouse and rat genes were identified using biomaRt R package to enable across-organism comparison. Spearman correlation (ρ) and a p-value <0.05 was considered significant and, hence, used to propose an experimental model as a potential PD study model. For each proposed model, their corresponding pathway scores were used to identify functional overlaps, depicting metabolic pathways in human PD that the animal PD model could be used to investigate.

## Author contributions

TÇ: Designed the study, oversaw the computational simulations and discussion of the results, wrote the manuscript, RO: Performed the computational simulations and discussion of the results, wrote the manuscript, EA: Designed the study, performed initial computational simulations and OB: Performed computational simulations.

## Conflict of interests

The authors declare availability of no competing interests.

## Funding

This study was financially supported by the Turkish Institutes of Health (TÜSEB) under grant number 2019-TA-01-3440.

## Materials and correspondence

All correspondences should be addressed to the corresponding author.

## Ethics approval

No ethical approval was required in this study as no human subjects or animals were used. The data used in this study are publicly available and no patient consent was required for their re-analysis.

## Data availability

All the data samples used in this study are publicly available in GEO ^53^, ArrayExpress ^20^, OMIM ^42^ and Human-GEM ^23^ data repositories. Where necessary accession numbers have been provided.

## Code availability

Base R scripts were used, and all packages used at each step are cited.

## Appendix: Supplementary information

### Supplement 1

**Fig. S1:** Cluster characterization using the pathway score approach. FC: gene fold change, PV: gene p-value.

**Fig. S2**: Clustering brain tissues using Pearson correlation as the distance metric. (a) human datasets (b) experimental model datasets. 729 genes, a subset of the 1525 genes used in clustering human datasets, were used in the clustering analysis of the experimental models.

**Fig. S3**: Correlation of gene expression between Substantia nigra and (a) DON and (b) ME. DON: Dorsal olivary nucleus, ME: Medulla. The points correspond to each gene whose scores were compared.

### Supplement 2

**Table S1**: A summary of the transcriptome data sets used in the study. This table provides information regarding the brain regions investigated by the original studies as well as the data collection platform used. The references provided are for the studies published in GEO associated with the studies. In some instances, the original data publishers in GEO are different from authors of the associated studies.

**Table S2**: Summary of novel metabolic genes identified in the Substantia nigra that had not been previously characterised in Parkinson’s disease.

## Notes

https://github.com/SysBioGTU/PDClustering

### Competing Interest Statement

The authors have declared no competing interest.

