## Supplementary figures and images for "Brain-wide transcriptome-based metabolic alterations in Parkinson’s disease: human inter-region and human-experimental model correlations"

### Supplementary Figure 1

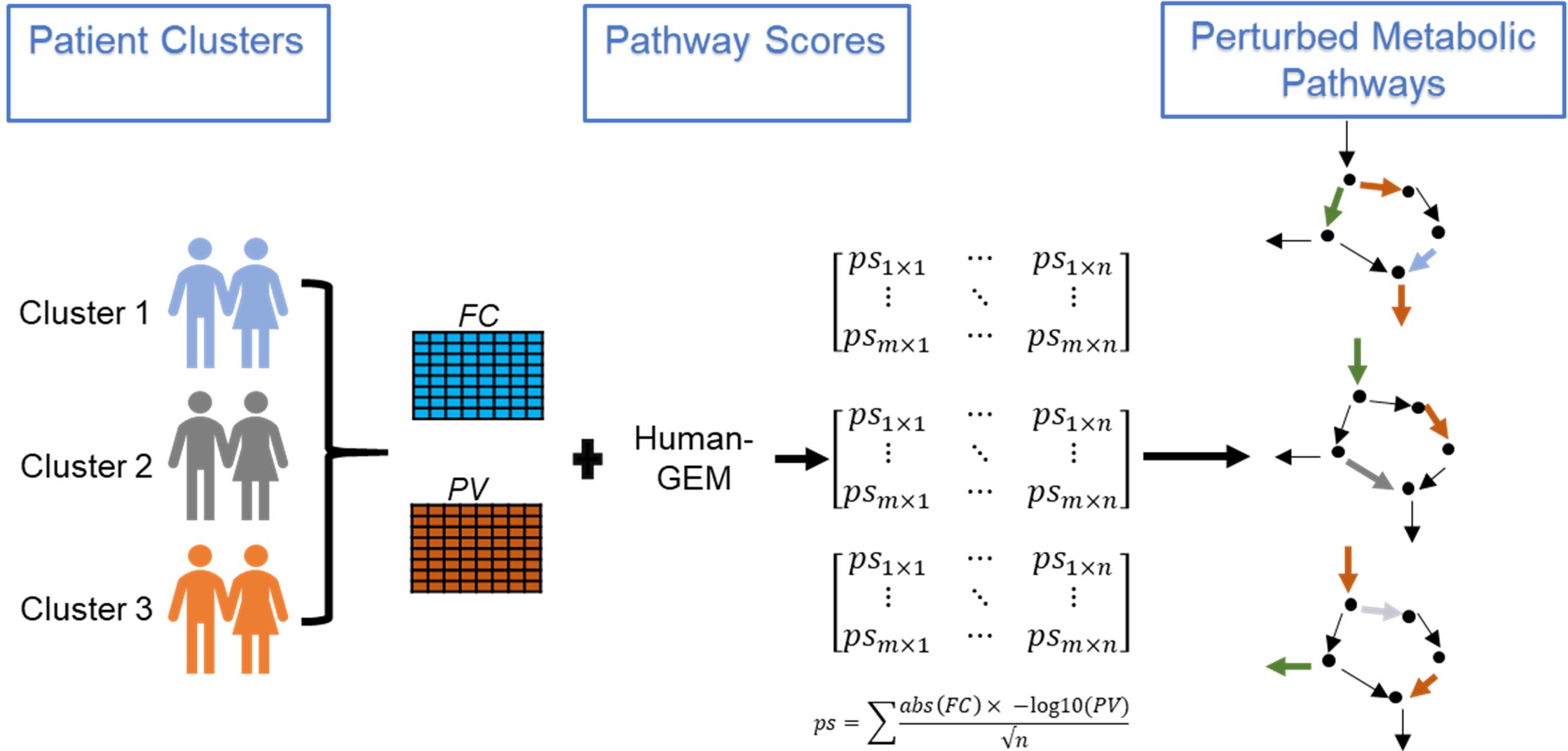

### Supplementary Figure 2

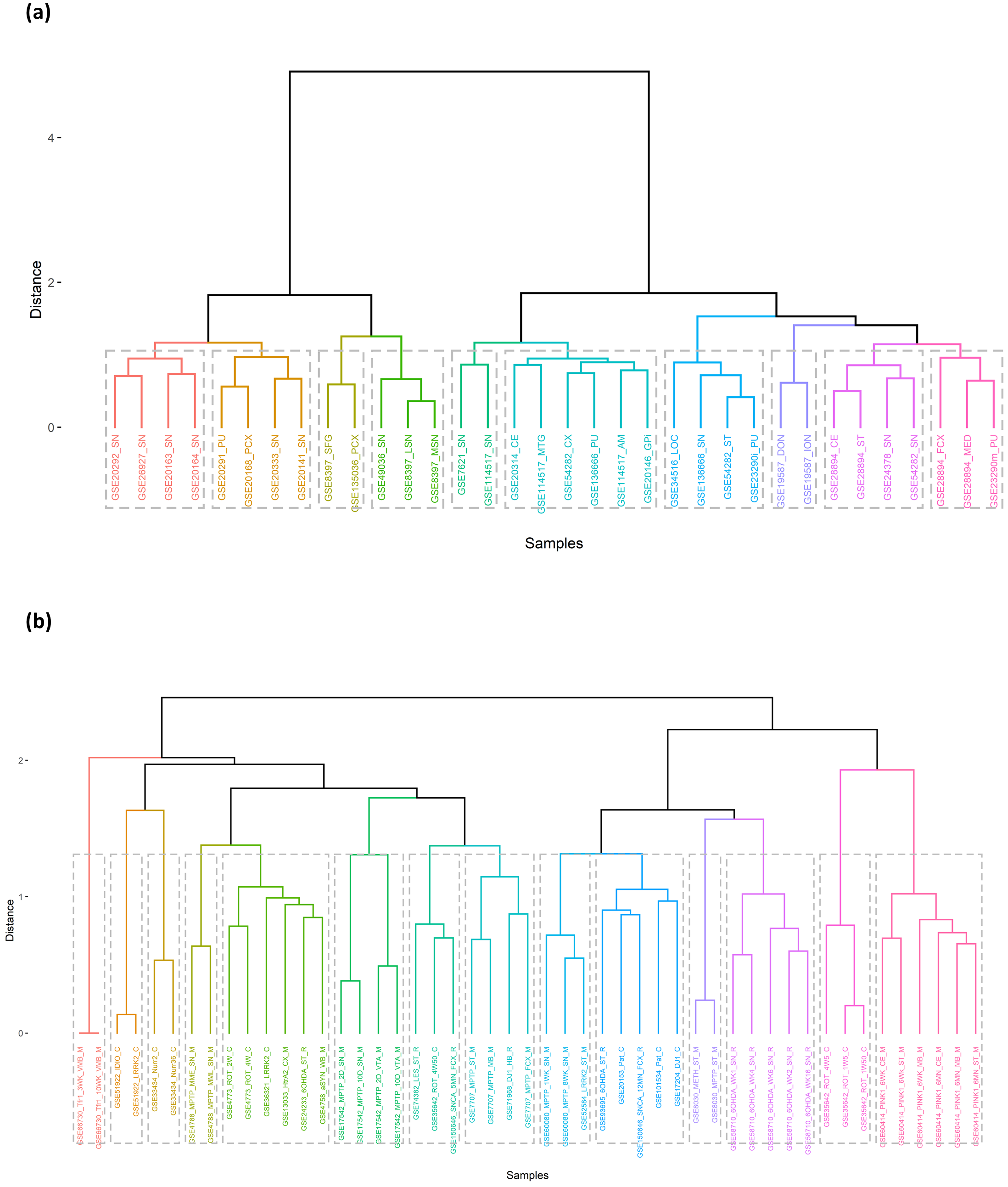

### Supplementary Figure 3

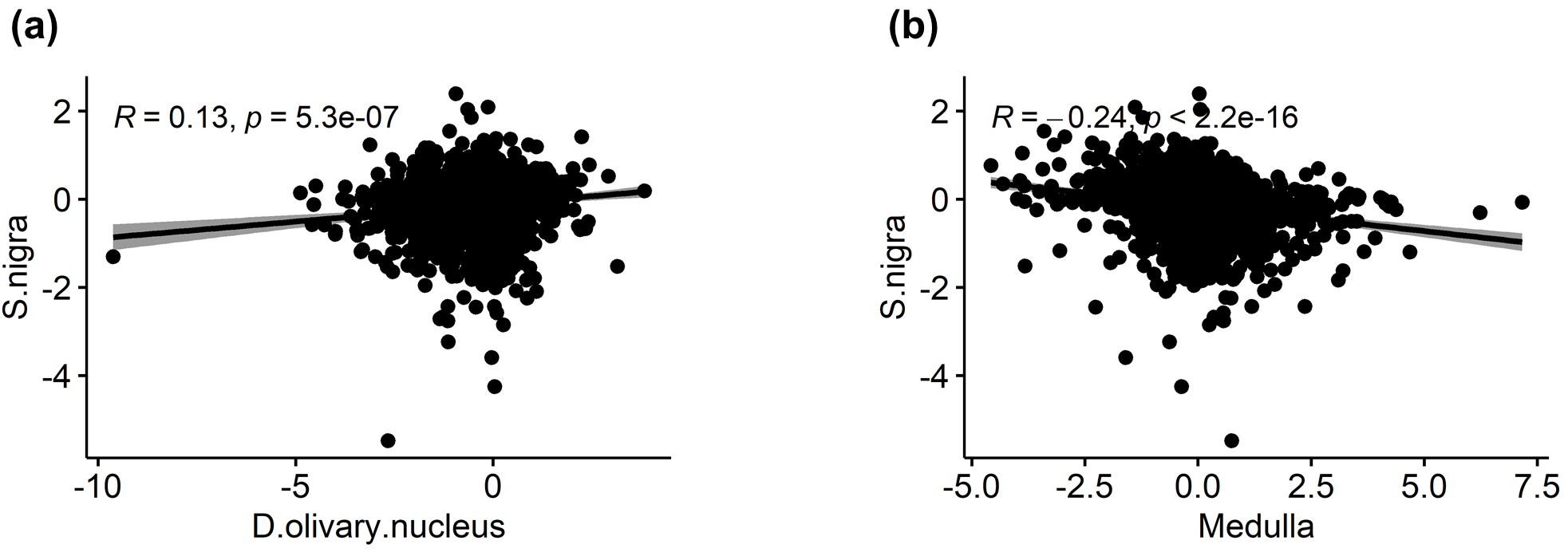
