## Supplementary Table 1 for "Brain-wide transcriptome-based metabolic alterations in Parkinson’s disease: human inter-region and human-experimental model correlations"

**Table S1**: A description of the transcriptome data sets used in the study. This table provides information regarding the brain regions investigated by the original studies as well as the data collection platform used. The references provided are for the studies published in GEO associated with the studies. In some instances, the original data publishers in GEO are different from authors of the associated studies.

| **Accession No.** | **Description** | **Platform** | **Reference** |
| --- | --- | --- | --- |
| GSE19587 | DON  ION | Affymetrix Human Genome U133A 2.0 Array | (Lewandowski et al., 2010) |
| GSE34516 | LOC | Affymetrix Human Exon 1.0 ST Array [transcript (gene) version] | (Botta-Orfila, Sànchez-Pla, et al., 2012) |
| GSE136666 | SN  PU | Illumina HiSeq 2000 (Homo sapiens) | (Xicoy et al., 2020) |
| GSE54282 | ST  SN  CO | Affymetrix Human Gene 1.0 ST Array [HuGene10stv1_Hs_ENTREZG_15.0.0] | (Riley et al., 2014) |
| GSE28894 | CO  ST  FCX  ME | Illumina humanRef-8 v2.0 expression beadchip | (Riley et al., 2014) |
| GSE24378 | SN | Affymetrix Human X3P Array | (Zheng et al., 2010) |
| GSE23290 | mPU  iPU | Affymetrix Human Exon 1.0 ST Array [transcript (gene) version] | (Botta-Orfila, Tolosa, et al., 2012) |
| GSE7621 | SN | Affymetrix Human Genome U133 Plus 2.0 Array | (Lesnick et al., 2007) |
| GSE114517 | SN  AM  MTG | Illumina NextSeq 500 (Homo sapiens) | (Simchovitz et al., 2020) |
| GSE20146 | GP | Affymetrix Human Genome U133 Plus 2.0 Array | (Zheng et al., 2010) |
| GSE20314 | CE | Affymetrix Human Genome U133A Array | (Zheng et al., 2010) |
| GSE20163 | SN | Affymetrix Human Genome U133A Array | (Zheng et al., 2010) |
| GSE20164 | SN | Affymetrix Human Genome U133A Array | (Zheng et al., 2010) |
| GSE20333 | SN | Affymetrix Human HG-Focus Target Array | (Zheng et al., 2010) |
| GSE20141 | SN | Affymetrix Human Genome U133 Plus 2.0 Array | (Zheng et al., 2010) |
| GSE20292 | SN | Affymetrix Human Genome U133A Array | (Zhang et al., 2005; Zheng et al., 2010) |
| GSE20291 | PU | Affymetrix Human Genome U133A Array | (Zhang et al., 2005; Zheng et al., 2010) |
| GSE20168 | PCX | Affymetrix Human Genome U133A Array | (Zhang et al., 2005; Zheng et al., 2010) |
| GSE8397 | SFG  LSN  MSN | Affymetrix Human Genome U133A Array | (Duke et al., 2007; Moran et al., 2006) |
| GSE135036 | PCX | Illumina NextSeq 500 (Homo sapiens) | (Li et al., 2020) |
| GSE26927 | SN | Illumina humanRef-8 v2.0 expression beadchip | (Durrenberger et al., 2012, 2015) |
| GSE49036 | SN | Affymetrix Human Genome U133 Plus 2.0 Array | (Dijkstra et al., 2015) |
| GSE4773 | SK-N-MC exposed to rotenone | Affymetrix Human Genome U133 Plus 2.0 Array | (Cabeza-Arvelaiz & Schiestl, 2012) |
| GSE17204 | DJ-1 mutant SH-SY5Y cell lines | Affymetrix Human Genome U133A 2.0 Array | (Foti et al., 2010) |
| GSE20153 | EBV transformed cell lines | Affymetrix Human Genome U133 Plus 2.0 Array | (Zheng et al., 2010) |
| GSE35642 | SK-N-MC exposed to rotenone | Affymetrix Human Genome U133A Array | (Cabeza-Arvelaiz & Schiestl, 2012) |
| GSE36321 | LRRK2 mutant neural stem cells | Affymetrix Human Genome U133A 2.0 Array | (Liu et al., 2012) |
| GSE101534 | LRRK2-G2019S mutant hNES cells | Affymetrix Human Gene 2.0 ST Array [transcript (gene) version] | (Arias-Fuenzalida et al., 2017; Bolognin et al., 2019) |
| GSE4788 | SN  exposed to MPTP | Affymetrix Murine Genome U74A Array | (Miller et al., 2004) |
| GSE4758 | WB  aSYN mouse line | Affymetrix Mouse Genome 430 2.0 Array | (De et al., 2010) |
| GSE7707 | PCX-ST-MB  exposed to MPTP | Affymetrix Mouse Genome 430 2.0 Array | (Williams, 2013) |
| GSE8030 | ST  Exposed to MPTP and METH | Affymetrix Mouse Expression 430A Array | (Chin et al., 2008) |
| GSE13033 | CX  Mutant HtrA2 | Affymetrix Mouse Genome 430 2.0 Array | (Moisoi et al., 2009) |
| GSE17542 | SN-VTA  Exposed to MPTP | Affymetrix Mouse Genome 430 2.0 Array | (Phani et al., 2010) |
| GSE60080 | ST  Exposed to MPTP-HCI, Saline | Affymetrix Mouse Gene 1.0 ST Array [transcript (gene) version] | (Nishimura et al., 2015) |
| GSE60414 | CE-MB-ST  Mutant Pink1 /A53T-SNCA | Affymetrix HT MG-430 PM Array Plate | (Gispert et al., 2015) |
| GSE66730 | VMB  Mutant Tfr1 | Affymetrix Mouse Genome 430 2.0 Array | (Matak et al., 2016; Sterky et al., 2012) |
| GSE52584 | ST  Mutant LRRK2 | Affymetrix Mouse Gene 1.0 ST Array [transcript (gene) version] | (Dorval et al., 2014) |
| GSE24233 | ST  Exposed to 6-OHDA/Saline | Illumina ratRef-12 v1.0 expression beadchip | (Cadet et al., 2010) |
| GSE58710 | SN  Exposed to 6-OHDA,  Vehicle | Affymetrix Rat Gene 1.0 ST Array [transcript (gene) version] | (Kanaan et al., 2015) |
| GSE74382 | DST  Exposed to saline | Affymetrix Rat Genome 230 2.0 Array | (Loiodice, 2016) |
| GSE93695 | ST  Exposed to L-DOPA | Affymetrix Rat Gene 2.0 ST Array [transcript (gene) version] | (Chen et al., 2017) |
| GSE71968 | HMB  Mutant DJ-1 | Illumina HiSeq 2000 (Rattus rattus) | (Hauser et al., 2017) |
| GSE150646 | PCX  Mutant SNCA | Illumina HiSeq 2500 (Rattus norvegicus) | (Hentrich et al., 2020) |

**SN**: Substantia nigra, **ST**: Striatum, **PU**: Putamen, **GP**: Globus pallidus, **AM**: Amygdala, **CE**: Cerebellum, **ME**: Medulla oblangata, **LC**: Locus coeruleus, **DON**: Dorsal motor nucleus of the vagus, **ION**: Inferior olivary nucleus, **PCX**: Prefrontal cortex, **CX**: Whole cortex, **MTG**: Medial temporal gyrus and **SFG**: Superior frontal gyrus, **WB:** Whole Brain, **MB:** Midbrain, **VTA**: Ventral Tegmental Area, V**MB:** Ventral Midbrain, **DST**: Dorsal Striatum, **HMB**: Hemibrain. **MPTP**: 1-methyl-4-phenyl-1,2,3,6-tetrahydropyridine, **6-OHDA**: 6-hydoxy-dopamine, **METH**: Methamphetamine, **L-DOPA**: levo-dopamine
