## Supplementary Table 2 for "Brain-wide transcriptome-based metabolic alterations in Parkinson’s disease: human inter-region and human-experimental model correlations"

**Table S2**: List of novel metabolic genes identified in the Substantia nigra that had not been previously characterised in Parkinson’s disease.

| **Gene ID** | **Gene Name** | **Enzyme** | **Protein Family** | **OMIM ID** | **Uniprot ID** | **Ontology** | **PMID** | **Genomic Location** | **RefSeq_MRNA** |
| --- | --- | --- | --- | --- | --- | --- | --- | --- | --- |
| ANXA3 | annexin A3 |  | PF00191 | 106490 | P12429 | GO:0005509 | 1830024 | 4q21.21 | NM_005139 |
| B3GAT1 | beta-1,3-glucuronyltransferase 1 | 2.4.1.135 | PF03360 | 151290 | Q9P2W7 | GO:0000139 | 8619474 | 11q25 | NM_054025,  NM_001367973,  NM_018644 |
| CAD | carbamoyl-phosphate synthetase 2, aspartate  transcarbamylase, and dihydroorotase | 3.5.2.3 | PF02787 | 114010 | F8VPD4 | GO:0001889 | 1979741 | 2p23.3 | NM_004341,  NM_001306079 |
| GK | glycerol kinase | 2.7.1.30 | PF00370 | 300474 | B4DH54 | GO:0004370 | 2081587 | Xp21.2 | NM_001128127,  NM_203391,  NM_001205019,  NM_000167 |
| GLTP | glycolipid transfer protein |  | PF08718 | 608949 | A0A024RBI7 | GO:0005515 | 2190982 | 12q24.11 | NM_016433 |
| GNE | glucosamine (UDP-N-acetyl)-2-epimerase/  N-acetylmannosamine kinase | 2.7.1.60 | PF00480 | 269921 | Q9Y223 | GO:0004553 | 2443758 | 9p13.3 | NM_001128227, NM_001190384, NM_001190388, NM_001374797,  NM_005476,  NM_001190383,  NM_001374798 |
| GUCY1B1 | guanylate cyclase 1 soluble subunit beta 1 | 4.6.1.2 | PF07700 | 139397 | Q02153 | GO:0004383 | 1352257 | 4q32.1 | NM_001291954, NM_001291952, NM_001291951, NM_001291955, NM_001291953,  NM_000857 |
| MAP4K2 | mitogen-activated protein kinase kinase kinase kinase 2 | 2.7.11.1 | PF00069 | 603166 | Q12851 | GO:0000139 | 7477268 | 11q13.1 | NM_001307990,  NM_004579 |
| PLOD3 | procollagen-lysine,2-oxoglutarate 5-dioxygenase 3 | 2.4.1.66 | PF03171 | 603066 | O60568 | GO:0001701 | 9582318 | 7q22.1 | NM_001084 |
| PLPP2 | phospholipid phosphatase 2 | 3.1.3.4 | PF01569 | 607126 | O43688 | GO:0005515 | 8889548 | 19p13.3 | NM_177543,  NM_003712,  NM_177526 |
| TM7SF2 | transmembrane 7 superfamily member 2 | 1.3.1.70 | PF01222 | 603414 | O76062 | GO:0005515 | 9286704 | 11q13.1 | NM_001277233,  NM_003273 |
